# Claustrum projections to the anterior cingulate modulate nociceptive and pain-associated behaviour

**DOI:** 10.1101/2024.01.18.576065

**Authors:** Christian A. Faig, Gloria H. K. Kim, Alison D. Do, Zoë Dworsky-Fried, Jesse Jackson, Anna M. W. Taylor

## Abstract

The anterior cingulate cortex (ACC) processes nociceptive information and pain unpleasantness. The claustrum is a subcortical region that provides robust feed-forward inhibition onto the ACC, suggesting this circuit could play a role in modulating pathological states such as pain. However, the function of this circuit in the context of acute and chronic inflammatory pain is unclear. Here, we show that claustrocingulate neurons exhibit a bimodal pattern of activation following acute pain but are suppressed during states of chronic inflammatory pain. Molecular lesion and chemogenetic suppression of claustrocingulate neurons increased acute nociception but interfered with pain learning. Activation of this pathway rescued mechanical allodynia associated with chronic pain. Together, these results suggest claustrocingulate neurons are a critical component of the pain neuromatrix, and dysregulation of this connection may contribute to chronic pain.

## 1. Introduction

The anterior cingulate cortex (ACC) is a critical node for pain processing^1-5^. It receives nociceptive information from regions such as the thalamus and amygdala, and projects to cortical and subcortical regions involved in pain processing, such as the striatum, periaqueductal grey, and frontal cortices^6,7^. In humans, activity in the ACC correlates with the reported perception of pain, rather than stimulus intensity^3,8^. However, activation of specific subpopulations of ACC neurons produces hypersensitivity to innocuous stimuli (i.e. allodynia)^9,10^. Chronic pain induces functional changes in central nervous system processing of pain, including ACC hyperexcitability^11-15^. Cingulotomy alleviates malignant, chronic low back, and neuropathic pain in humans^16,17^. Similarly, inhibition of ACC activity reverses mechanical allodynia in animal models of persistent and neuropathic pain^1,9,18,19^. These experiments highlight the role of the ACC in encoding of both sensory and affective components of acute and chronic pain.

A less well studied projection into the ACC arises from the claustrum^20,21^. The claustrum is a small subcortical structure situated between the insula and striatum^21-23^. Retroand anterograde neuronal tracing has unveiled dense reciprocal connections with several cortical regions^21-25^. The densest of these claustrum projections target frontal cortical structures, including the ACC. Optogenetic stimulation of claustrum projection neurons activates cortical parvalbumin (PV+) interneurons that generates feed-forward inhibition onto excitatory cortical networks^27,28^. Activation of ACC PV+ interneurons rescues mechanical nociceptive thresholds in an animal model of chronic pain^9^, suggesting claustrum inputs may function to attenuate pain processing. Despite prior work characterizing claustrocortical anatomy and physiology, the functional role of specific claustrum connections in the context of acute and chronic pain remains elusive.

Previous work has shown claustrocingulate projections display altered activity in prolonged pain^29-31^. However, it remains unclear if and how the claustrum participates in nociceptive processing and high-order pain behaviors. In this study, we sought to define the function of claustrocingulate projections in acute and chronic pain. We found that claustrum activity was enhanced after a painful stimulus, and this response was attenuated in chronic inflammatory pain. Selective inhibition of claustrocingulate projection neurons enhanced acute nociception, but blocked pain learning. Inversely, chemogenetic claustrocingulate activation had no effect on basal nociception but rescued mechanical allodynia induced by persistent and chronic inflammation. Together, these results implicate claustrocingulate projections in nociceptive processing and identify a novel circuit dysfunction in chronic pain.

## 2. Methods

### 2.1 Animals

Adult male and female (8-10 weeks) C57BL/6J mice were obtained from Jackson Laboratories (Bar Harbor ME). Mice were housed in ventilated plastic cages in groups of two to four with standard bedding, free access to standard rodent chow and were kept on a 12-hour light/dark cycle with lights on at 7:00 am. All animal care and experiments were done in accordance with Canadian Council on Animal Care Guidelines and Policies and approved by the University of Alberta Health Sciences Animal Care and Use Committee.

### 2.2 Drugs and viruses

Clozapine n-oxide (CNO, CederLane BML-NS105-0025) was dissolved in saline (0.9% sodium chloride) and 3% DMSO and was given by inter-peritoneal (IP) injection at 2.5 mg/kg. Cholera Toxin subunit-B with an Alexa Fluor-647 (CTB, ThermoFischer C34778) conjugate was prepared prior to injection by dissolving 100 µg in 50 µl 1x PBS and stored at 4°C^32^. Adenosine associated viruses (AAV) were acquired from Addgene and stored at -80°C in 3 µl aliquots. Prior to injection, viral aliquots were diluted 1:1 with sterile 1x PBS. pENN.AAV.hSyn.HI.eGFPCre.WPRE.SV40 (Addgene viral prep # 105540-AAVrg; http://n2t.net/addgene:105540; RRID:Addgene_105540) was injected into the ACC (200 nl). pAAV-flex-taCasp3-TEVp (Addgene viral prep # 45580-AAV5; http://n2t.net/addgene:45580; RRID:Addgene_45580)^33^, pGP-AAV-syn-FLEX-jGCaMP7f-WPRE (Addgene viral prep # 104492-AAV1; http://n2t.net/addgene:104492; RRID:Addgene_104492)^34^, pAAV-hSyn-DIO-hM3D(Gq)mCherry (Addgene viral prep # 44361-AAV5 ; http://n2t.net/addgene:44361; RRID:Addgene_44361)^35^, or pAAV-hSyn-DIO-hM4D(Gi)-mCherry (Addgene viral prep # 44362AAV5; http://n2t.net/addgene:44362; RRID:Addgene_44362)^35^ was injected into the claustrum (200 nl).

### 2.3 Von Frey

Mechanical withdrawal thresholds were measured by the von Frey assay. Animals were repeatedly habituated to a 7.5x10x7.5 cm plexiglass box on a suspended mesh platform and allowed to explore for 30 minutes. On test days, the up-down method was used to determine the 50% withdrawal threshold^36^. Briefly, filaments of various force (0.04-4g) were applied to both hind paws and positive and negative responses were recorded. The 50% withdrawal threshold was calculated, as previously described^37^. An alternative assay of 10 stimuli 10 seconds apart with either an innocuous filament (0.4g) or noxious filament (1.4 or 2g) was performed on mice 90-120 minutes before euthanasia. The number of vigorous paw withdrawals were recorded. Noxious and innocuous filament strength was selected based on historical 50% threshold averages. The noxious filament (1.4g) produces nociceptive responses in 80% of control mice. The innocuous filament (0.4g) produces nociceptive responses in less than 20% of mice.

### 2.4 Tail withdrawal Assay

Thermal withdrawal thresholds were measured by tail withdrawal assay. Mice were habituated to acute restraint stress. Animals were gently restrained in a soft conical tube and the distal 2.5 cm of their tail was immersed in 49°C water. The time to withdrawal was measured, and a cutoff latency of 15 seconds was set to prevent tissue damage. Three measurements were collected 15 minutes apart and later averaged. Following a subcutaneous injection of CFA in the left hind paw, measurements were collected 3and 14-days post injection.

### 2.5 Conditioned place aversion and acetic acid writhing

Acetic acid conditioned place aversion (CPA) was conducted using a biased 2-chamber apparatus. The 2 chambers (24 × 24.5 × 28 cm) are distinguished by spot or stripe patterns on the wall and separated by a guillotine door. Mice were placed in the CPA apparatus and allowed to explore freely for 20 minutes. Time spent in each chamber was recorded by overhead infrared cameras paired with a computer running behavioural tracking software (EthoVision, Noldus). The acetic acid paired chamber was assigned to the least preferred chamber. Mice were conditioned with a trial of acetic acid (1% acetic acid, i.p.) a trial of saline vehicle (0.9% NaCl, i.p.) for 4 trials of each condition alternating over 8 days. Animals were confined to the paired chamber for 20 minutes following injection before being returned to the home cage. On the final testing day, mice were not injected but were allowed to explore both chambers for 20 minutes. The amount of time spent in each chamber was assessed.

### 2.6. Inflammatory pain model

Mice were tested for baseline thresholds with the Von Frey and tail withdrawal assays the day prior to hind paw injection. They were then restrained before receiving a subcutaneous injection of 50 µl complete Freund’s adjuvant (CFA; 1mg/mL Sigma) or saline to the left hind-paw using a 27g needle. Behavioral testing was then repeated at 3-, 14-, and 21-days post injection.

### 2.7 Stereotaxic Injection

Mice were initially anesthetized at 4% isoflurane and maintained at 1.5-2.5% before being affixed to a stereotaxic frame. Body temperature was maintained at 37°C with an electric heat pad. Bupivacaine was administered locally, and an incision was made along the midline to expose bregma and injection sites. The skull was leveled using bregma and lambda prior to manual craniotomy of the injection sites using a 400 µm dental drill bit. Mediolateral and anteroposterior stereotaxic coordinates were measured from bregma and dorsoventral measurements were made from brain surface. All injections were performed bilaterally, targeting either the anterior cingulate cortex (A/P: +0.7, M/L: ±0.5, D/V: -0.75) or claustrum (A/P: +1.25, M/L: ±2.5, D/V: -2.5). Pulled pipettes (10 µm) were back filled with mineral oil prior to virus or tracer loading. Loaded pipettes were lowered to each coordinate at a rate 1mm per minute and 200 nl of tracer was injected at a rate of 25-50 nl per minute via pressure injection. Pipettes rested for 10 minutes before retraction. Mice were administered carprofen 24 hours prior to and 72 hours following operation via ad-libitum water to achieve a dose of 5 mg/kg.

### 2.8 Optic cannula implantation and headplating

Two weeks after stereotaxic injection, mice were planted with a headplate and optic fiber cannula using the general surgical procedure above. The skull was exposed and cleaned before light abrasion with a scalpel to improve adhesion. A drop of superglue was placed on the skull and a custom stainless steel headplate with a circular window was lowered onto the glue as it dried. After manual craniotomy, a 400 µm core optic fiber cannula (RWD, cat No. R-FOCBL400C-39NA) was lowered into the brain at a rate of 1 mm per minute until it was just above the injection site (A/P: +1.25, M/L: ±2.5, D/V: -2.45). The cannula and the headplate were then fixed in place using a strong dental cement (C & B Metabond). Mice were allowed to recover as above over 2 weeks before testing for positive signal.

### 2.9 Fiber Photometry

Fiber photometry recordings were acquired with a Doric basic fiber photometry system (Doric Lenses). Continuous light was produced from a 470 nm blue-light LED to excite GCaMP at 10-100 µW using a generated lock in frequency. Emitted light was collected and filtered by a Doric mini cube (FMC6_IE(400-410)_E1(460–490)_F1(500–540)_E2(555–570)_F2(580–680)_S) and transmitted as a voltage signal registered by Doric Neuroscience Studios. Recording files were processed and analyzed in Matlab 2022b, fitting an exponential decay to correct for photobleaching and generating a z-score to normalized data for ease of cross-animal comparisons.

Mice were habituated to head fixation over one week. Baseline recordings of claustrocingulate activity following a series of 5 von Frey stimulations with an innocuous and a noxious filament, or 6 trials of a tail withdrawal assay on separate recording days. The onset of stimulus and mouse response were manually recorded by the experimenter using a voltage pulse to generate time-locked digital currents within Doric Neuroscience Studios for analysis of stimulus evoked changes in activity. Mice then received a hindpaw injection of CFA while connected to the apparatus to assess activity changes caused by a robust nociceptive event. Recordings of acute nociceptive stimuli were also acquired at 3- and 14-days post CFA injection. Average activity across 4 10-minute epochs was assessed and compared: pre-behavior, during behavior, 0-10 minutes post behavior, 10-20 minutes post behavior. Stimulus evoked activity was assessed by averaging the recorded fluorescence 10 seconds before and after each repeated stimulus and plotting the averaged z-score over time.

### 2.10 Perfusion and Tissue Sectioning

Surgical anesthesia was induced by injection of sodium pentobarbitol (0.1mL, i.p., Euthanyl). Mice were then flushed with ice-cold 4% paraformaldehyde solution via intracardial perfusion. Brains were then extracted and bathed in 4% paraformaldehyde for 48 hours at 4°C. They were then cryopreserved in 30% sucrose, frozen in Optimal Cutting Temperature (OCT Tissue-Tek, Sakura, 4583) and sectioned into 20 µm slices on a cryostat before being mounted onto slides.

### 2.11 Immunohistochemistry

Slides were washed in 0.1M phosphate buffered saline (PBS) and then blocked with 10% normal donkey serum for 1 hour at room temperature. Sections were incubated with primary antibody rabbit anti-c-Fos (1:3000, New England Biolabs cat. No. 2250S) for 24 hours at room temperature. The slides were then incubated with secondary antibody donkey anti-rabbit Alexa Fluor 488 (1:200) or donkey anti-rabbit Alexa Fluor 594 (1:200) at room temperature for 45 minutes. The slices were washed twice with 0.05% tween in PBS and once with PBS before adding mounting medium and lowering a coverslip. For somatostatin immunohistochemistry, slices were incubated in rat anti-somatostatin (1:250, Millipore cat. No. MAB354, RRID:AB_2255365) at 4°C for 48 hours before the addition of primary antibody rabbit anti-c-Fos (1:3000), they were then incubated a further 24 hours at room temperature. For parvalbumin/cFos immunohistochemistry slices were incubated in goat anti-parvalbumin (1:2000, Swant Antibodies cat. No. PVG213) and rabbit anti-c-Fos (1:3000) for 24 hours at room temperature. Appropriate secondary antibodies were used to visualize each stain including goat anti-rat Alexa fluor 555 (1:200) and donkey anti-goat Alexa fluor 555 (1:200).

### 2.12 Statistical analysis

All behavioural and tissue analysis was performed by experimenters blinded to treatment. Statistical analysis was performed using GraphPad Prism software version 10.0.1 All data was tested for normality and parametric or non-parametric tests were used accordingly. Paired or unpaired t-tests were used to compare single treatment effects against a control. Multiple group means were analyzed using 1o-r 2-way analysis of variance (ANOVA) followed by Sidak post-hoc analysis to correct for multiple comparisons. A Welsh ANOVA was used to correct for unequal variances. Within-animal comparisons across multiple timepoints of CFA inflammation were analyzed using a mixed effects analysis with Sidak post-hoc test. Correlations were calculated using Pearson’s r. All data, aside from correlations, are represented as mean ± S.E.M. Significance threshold was set at p < 0.05 for all experiments.

## 3. Results

### 3.1 Acute and persistent pain activate the claustrum

To assess the activity of claustrocingulate projections in acute nociception, we injected Alexa Fluor 647 conjugated cholera toxin b subunit (CTB) into the ACC of male and female C57BL6/J mice. A noxious (1.4 g calibrated filament) or innocuous (0.4 g) mechanical stimulus was applied 10 times to each hind paw (**Figure 1A**). Mice were then euthanized and immunohistochemistry against c-Fos was used as a marker for neuronal activation^38^. Following a noxious stimulus, c-Fos expression in the claustrum was significantly increased (**Figure 1B, C**). When we restricted our analysis to c-Fos expression in CTB+ cells (projecting to the ACC), we also noted a significant increase of c-Fos expression, indicating robust pain-evoked activation of claustrocingulate neurons. No significant differences between males and females were found, and so sexes were pooled for all analyses. Application of innocuous mechanical stimuli failed to evoke any c-Fos expression in the claustrum (**Figure 1B, C**). Neither noxious or innocuous stimuli evoked c-Fos expression in PV+ interneurons in the claustrum, and c-Fos expression in SOM+ interneurons was similar between innocuous and noxious stimulation (**Supplemental Figure 1B**). To test whether persistent or chronic inflammatory pain induce similar activity, mice received a hind-paw injection of saline or CFA^39^. This produced a robust and long-lasting hind paw mechanical allodynia (**Figure 1D**). Hindpaw inflammation increased c-Fos expression in the claustrum similarly to acute noxious stimuli 72 hours following injection (**Figure 1E**). However, at 14 days following CFA, claustrocingulate c-Fos expression was not elevated (**Figure 1F**), even following noxious stimuli. These data suggest that claustrocingulate cells are activated by painful stimuli, but that activity in these neurons is suppressed in chronic pain.

**Fig 1.**
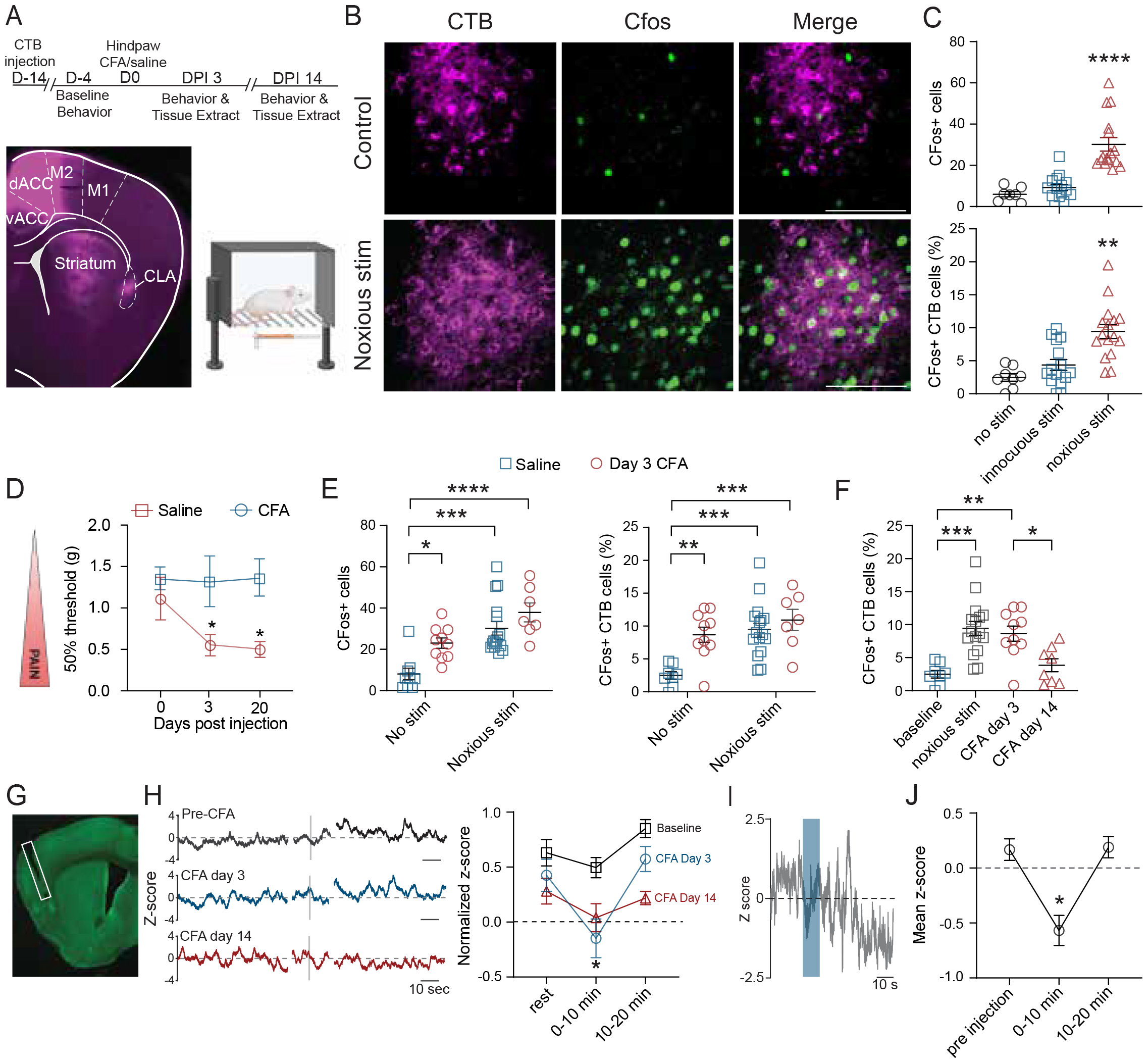
Acute and persistent pain activate the claustrum. (A) Timeline of stereotaxic injection and pain behaviour before euthanasia. An example image of a CTB injection with overlaid anatomy. A Biorender illustration of a mouse during the von Frey assay. (B) Example images of CTB and c-Fos co-labelling in a naïve mouse and after acute stimulation (scalebar = 100µm). (C) Effect of innocuous touch and noxious poke on c-Fos expression in the claustrum compared to naïve animals (n = 8-16, male and female mice) (W(2.000, 22.24) = 23.10, p < 0.0001; W(2.000, 23.91) = 16.51, p < 0.0001). (D) 50% paw withdrawal thresholds following a hind-paw injection of saline or CFA (n = 8, male and female mice) (effect of CFA: F(1,38)=12.0, p=0.0013). (E) Persistent inflammation and acute noxious stimulus similarly increased c-Fos expression in the claustrum (n = 8-16, male and female mice) (effect of stimulus: F(1, 38) = 27.43, p < 0.0001; effect of CFA: F(1, 38) = 10.30, p = 0.0027; effect of stimulus: F(1, 37) = 14.21, p = 0.0006, effect of CFA: F(1, 37) = 9.677, p = 0.0036. (F) Noxious stimulus failed to evoke c-Fos expression in the claustrum after chronic inflammatory pain (n = 8-16, male and female mice) (F(3, 38) = 10.15, p < 0.0001). (G) Example image of a cannula implantation over the claustrum. (H) Example trace of claustrum activity before, during and 20 minutes after nociceptive stimulation pre-CFA, 3- and 14-days post injection. Mean normalized z-score of claustrum activity 0-10 minutes and 10-20 minutes after tail flick (n = 7, male mice) (effect of timepoint: F(1.174, 8.221) = 8.893, p = 0.0147; effect of CFA: F(1.653, 11.57) = 16.40, p = 0.0006). (I) Example trace of claustrum activity during a hind-paw injection of CFA. (J) Mean z-score of claustrum activity 0-10 minutes and 10-20 minutes after CFA injection (n = 5, male mice) (F(1.510, 7.551) = 11.12, p = 0.0075). *p < 0.05, **p < 0.01, ***p < 0.001, ****p < 0.00001. Bars or points with error bars represent mean ± SEM. Data was analyzed by one-way or repeated measures ANOVA with Sidak’s post hoc test or a Welch ANOVA.

### 3.2 Claustrocingulate neurons respond to nociceptive stimuli in a temporally specific manner

To assess real-time claustrum activity and compliment c-Fos experiments, we used calcium imaging fibre photometry to study claustrocingulate cells during and after application of nociceptive stimuli in acute and chronic pain states. A dual injection strategy was used to express the calcium sensor (GCaMP7f) in claustrocingulate neurons, via the injection of AAVretro-Syn-Cre into the ACC and AAV1-FLEX-jGCaMP7f into the claustrum. Claustrum activity was recorded before, during, and after the application of noxious stimuli. Acute noxious thermal stimuli were applied to the tail during the pre-injection day, and 3 and 14 days later. Despite eliciting a tail flick response at each noxious stimulus, claustrocingulate activity did not elicit a change during the time of the stimulation (**Figure 1H**). However, the mean activity of the claustrum decreased immediately after the barrage of nociceptive stimuli, before increasing 10 to 20 minutes after the final stimulus (**Figure 1H**). We also observed that immediate injection of hindpaw CFA induced a robust decrease in claustrocingulate activity that recovered by 20 minutes post injection (**Figure 1I-J**). At 3 days post injection, there was an exaggerated decrease in claustrum activity following nociceptive stimuli before a similar increase 10 to 20 minutes after the final stimulus. However, at 14 days after CFA noxious stimulus was no longer able to evoke any change in claustrum activity (**Figure 1H**). This demonstrates that claustrocingulate activity is recruited in the period following a painful stimulus. However, the transition from acute to chronic pain is accompanied by reduced pain-evoked claustrum activity.

### 3.3 Selective claustrum inhibition promotes pain hypersensitivity

To determine the role of claustrocingulate projections in pain processing we selectively ablated claustrocingulate projections using viral directed Caspase-3 expression (**Figure 2A**). Claustrocingulate neurons were directly lesioned by injection of pAAV-flex-taCasp3-TEVp into the claustrum and a retrograde cre AAV into the AAC, as above. Mechanical thresholds were significantly reduced following claustrocingulate lesion indicating mechanical hypersensitivity (**Figure 2B**). Acute thermal nociception was not significantly affected by claustrocingulate lesion; however, the onset of CFA-induced systemic thermal hyperalgesia was accelerated (**Supplemental Figure 3A**). Reflexive and attending nociceptive behaviours in response to mechanical stimulus were recorded and analysed as a measure of higher order pain behaviours. In claustrocingulate lesioned mice, immediate reflexive nociceptive responses (paw flick, toe spread) were significantly reduced, though prolonged attending behaviours (paw guarding, grooming or shaking) were unchanged (**Figure 2B**). In control mice, heightened reflexive behaviours correlated with heightened attending behaviours, identifying a signature of pain behaviours in control mice that was not exhibited in claustrocingulate lesioned mice (**Figure 2B**). We then assessed spontaneous pain behaviour following an intraperitoneal injection of 1% acetic acid. Claustrocingulate lesioned mice showed increased writhing behaviour while overall locomotor activity was unchanged (**Figure 2C**). To determine if claustrocingulate lesioned mice could learn the association between spatial context and pain, we assessed pain learning to 1% acetic acid in a conditioned place aversion paradigm (**Figure 2D**). Control mice displayed a robust aversion to the acetic acid paired chamber, whereas mice with a claustrocortical lesion mice did not (**Figure 2D**). This suggests loss of claustrocingulate neurons impair pain avoidance learning.

**Fig 2.**
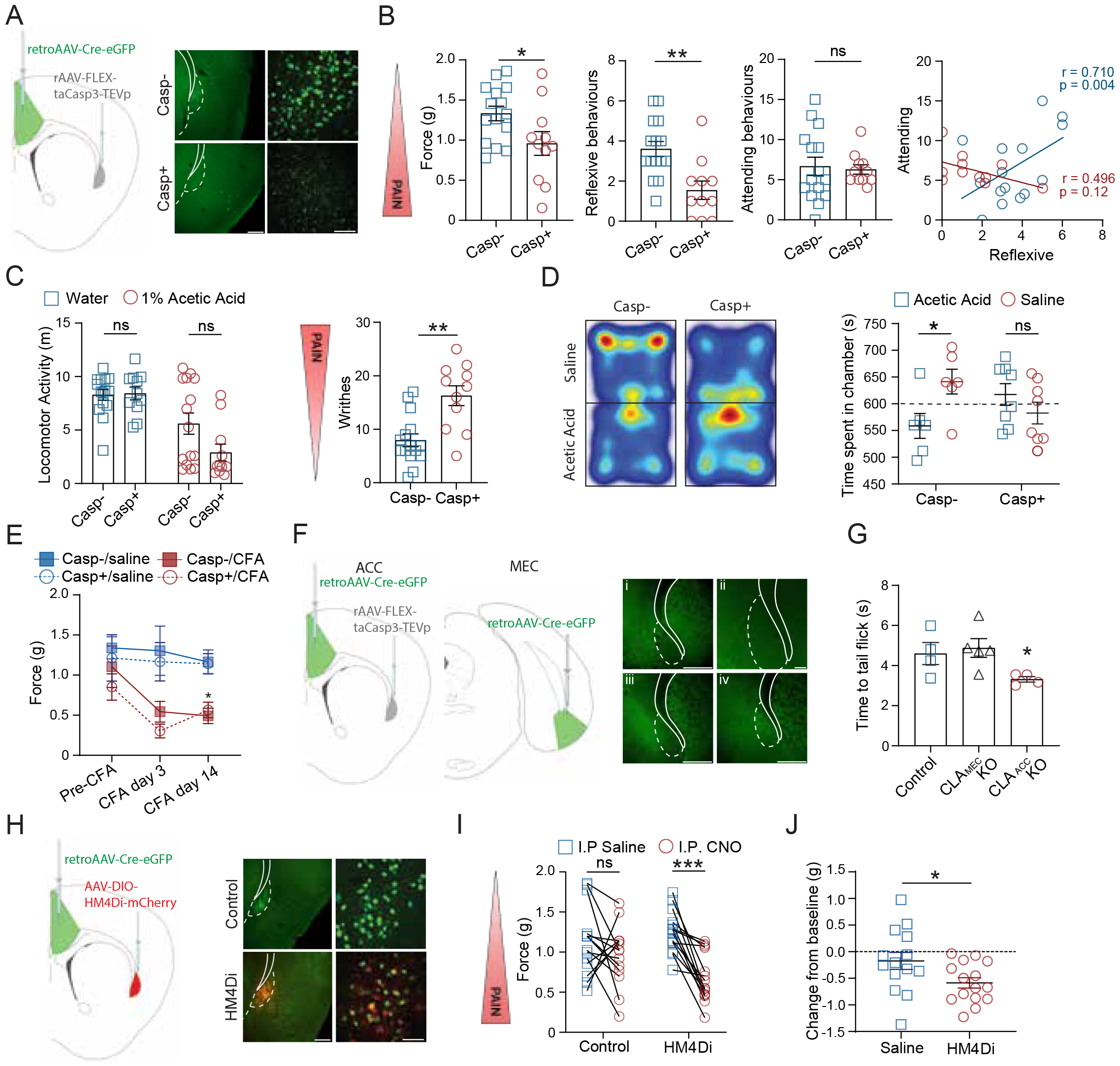
Selective claustrum inhibition induces pain hypersensitivity. (A) Schematic of virus directed claustrocingulate lesion and example images at 10x (scalebar = 200 µm) and 40x (scalebar = 50 µm). (B) Caspase positive mice show decreased paw withdrawal thresholds and reflexive nocifensive behaviors compared to controls, disrupting the correlation between reflexive, and attending behaviors (n = 11-15) (t(24) = 2.271, p = 0.0324; t(24) = 3.505, p = 0.0018; t(24) = 0.2755, p = 0.7853; r(14) = 0.7101, p = 0.0044; r(10) = -0.4969, p = 0.12). (C) Caspase positive mice do not show a difference in locomotor activity but show increased writhe behavior after an i.p. injection of 1% acetic acid (n = 11-15) (effect of caspase: F(1, 48) = 2.771, p = 0.1025, effect of acetic acid: F(1, 48) = 28.14, p < 0.0001; t(10) = 3.621, p = 0.0047). (D) Representative heatmap and quantification of time spent in 1% acetic acid paired chamber in a CPA paradigm (n = 6-8). Caspase mice failed to learn an aversion(t(5) = 4.053, p = 0.0154; t(7) = 0.8623, p = 0.4171). (E) 50% paw withdrawal thresholds of caspase positive mice are not different from control following hind-paw injection of CFA (n = 5-8) (effect of time: F(1.464, 47.58) = 2.873, p = 0.0815; effect of caspase: F(1, 65) = 1.034, p = 0.3129; effect of CFA: F(1, 65) = 25.12, p < 0.0001). (F) Schematic of virus directed CLA-ACC and CLA-MEC lesion with example images of CLA-ACC lesion at 10x (i, scalebar = 500 µm) and 20x (ii, scalebar = 100 µm), CLA-MEC lesion at 10x (iii) and control at 10x (iv). (G) tail withdrawal latency was decreased by CLA-ACC lesion but not CLA-MEC lesion (n = 4-5) (H(2) = 6.382, p = 0.0288). (H) Schematic of virus directed HM4Di expression and example images at 10x (scalebar = 200 µm) and 40x (scalebar = 50 µm). (I) Nociceptive thresholds after saline or CNO injection in control and hM4Di expressing mice and the differential (n = 15-16) (effect of CNO: F(1, 27) = 17.04, p = 0.0003; effect of HM4Di: F(1, 27) = 0.9602, p = 0.3358; CNO x HM4Di: F(1, 27) = 5.021, p = 0.0335). (J) CNO injection significantly decreased paw withdrawal thresholds in hM4Di expressing but not control mice (t(27) = 2.241, p = 0.0335). *p < 0.05, **p < 0.01. Bars or points with error bars represent mean ± SEM. Data was analyzed by paired and unpaired t tests, or repeated measures ANOVA with Sidak’s post hoc test. Correlations were assessed by Pearson’s r. Data was collected in male mice.

To assess the impact of claustrocingulate lesions on chronic pain, a separate group of mice received selective lesion of claustrocingulate neurons followed by a hind-paw CFA injection. While, lesioning the claustrocingulate neurons lowered mechanical nociceptive thresholds in pain naïve mice, it did not exacerbate the mechanical allodynia observed following hindpaw CFA injection (**Figure 2E**). We next assessed the effect of lesioning other claustrocortical pathways on nociceptive thresholds. Selective lesion of claustrum projections to the medial entorhinal cortex did not affect nociceptive thresholds, indicating that the impact on nociceptive behaviors is specific to claustrocingulate projections (**Figure 2F-G**).

Next, we determined how acute suppression of claustrocingulate neurons impacts nociceptive thresholds. Claustrocingulate neurons were targeted by injection of AAV5-FLEX-HM4Di-mcherry to the claustrum and retrograde cre AAV into the ACC, as above. Mechanical thresholds were tested following an injection of saline or clozapine-N-oxide (CNO) to suppress claustocingulate activity. HM4Di expressing mice had significantly reduced thresholds compared to control (not expressing the HM4Di receptor) mice that received CNO (Fig 2I-J). This result mirrors the effects of claustrocingulate ablation and suggests inhibition of claustrocingulate neurons lower nociceptive thresholds while interfering with higher order pain behaviours.

### 3.4 Selective claustrum activation rescues mechanical allodynia

Thus far, our data suggests claustrocingulate neurons are hypoactive during chronic pain and experimentally reducing claustrocingulate activity lowers nociceptive thresholds. Therefore, we hypothesized that activation of claustrocingulate neurons may reduce the expression of pain behavior in acute or chronic pain. We activated claustrocingulate neurons using virus-directed expression of AAV5-FLEX-HM3Dq-mcherry to examine the impacts on pain and nociception. Baseline mechanical and thermal nociceptive thresholds, reflexive, and attending pain behaviors were unaltered by claustrocingulate activation (**Figure 3B, supplemental Figure 3B**). Therefore, claustrocingulate activation has no discernible impact on baseline acute nociceptive behaviors. We then injected CFA to the hind paw to study claustrocingulate activation during chronic pain. Activating claustrocingulate neurons significantly increased mechanical thresholds in CFA injected mice at both the 3- and 14day timepoint (**Figure 3D-E**). In control mice that did not express HM3Dq, CNO did not significantly alter CFA induced mechanical allodynia. Therefore, claustrum activation does not impact acute nociception, but rescues the allodynia associated with chronic inflammatory pain.

**Fig 3.**
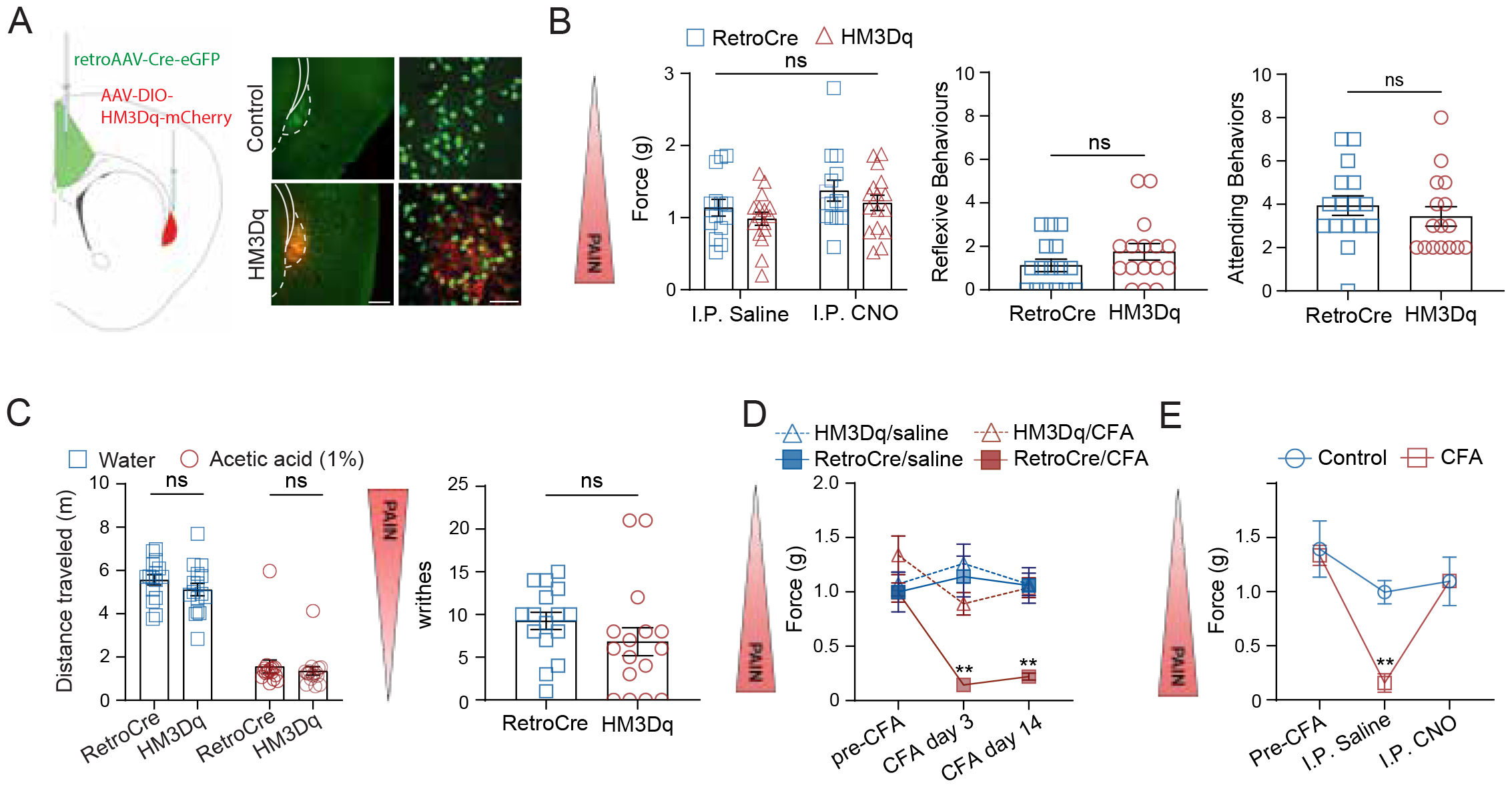
Selective claustrum activation does not affect basal nociception but does rescue mechanical allodynia. (A) Schematic of virus directed HM3Dq expression and example images at 10x (scalebar=200 µm) and 40x (scalebar = 50 µm). (B) Mechanical nociceptive thresholds following a saline or CNO injection in control and hM3Dq expressing mice (n = 15-16)(effect of drug: F(1, 56) = 1.986, p = 0.1643; effect of HM3Dq: F(1, 56) = 3.972, p = 0.0511). No effect of drug or virus was found on nociception, or nocifensive behaviours (t(30) = 1.309, p = 0.2006; t(30) = 0.7870, p = 0.4375). (C) Acute claustrum activation does not change locomotion or writhing behaviours following i.p. injection of 1% acetic acid (n = 15-16) (effect of HM3Dq: F(1, 30) = 1.630, p = 0.2115; effect of acetic acid: F(1, 30) = 222.7, p < 0.0001, t (30) = 1.265, p = 0.2157). (D) Claustrum activation significantly increases 50% paw withdrawal thresholds in mice that received a hind-paw injection of CFA compared to virus controls (n = 7-8) (effect of time: F(1.867, 50.42) = 6.718, p = 0.0031; effect of HM3Dq: F (1, 27) = 14.61, p = 0.0007; effect of CFA: F (1, 27) = 12.82, p = 0.0013; HM3Dq x CFA: F(1, 27) = 9.508, p = 0.0047). (E) Activation of claustrocingulate projections 3 days post CFA injection rescues nociception in hypersensitive mice (n = 3) (effect of CFA: F(1, 4) = 7.908, p = 0.0482; effect of CNO: F(1, 4) = 19.15, p = 0.0119; effect of CNO x CFA: F(1, 4) = 12.52, p = 0.0241). *p < 0.05, **p < 0.01. Data was analyzed by paired and unpaired t tests, or repeated measures ANOVA with Sidak’s post hoc test or Kruskal-Wallis test with Sidak’s post hoc test. Data was collected in male mice.

## 4. Discussion

The ACC plays many functions in the sensory and emotional processing of pain^2,3,40,41^. Distinct circuits within the ACC likely modulate these diverse functions through connections to varied subcortical and cortical connections^42,43^. Previous research has demonstrated that hyperactivity of the ACC is a hallmark of chronic pain and correlates with the onset of hyperalgesia and allodynia^12,14,15,^ Inhibition or lesion of the ACC reverses pain hypersensitivity in animal models of chronic pain and human cingulotomy reduces pain in various disease states^9,15-17^. Together, this suggests that ACC hyperactivity is a key contributing factor in chronic pain and amelioration of this hyperactivity could provide therapeutic benefits to chronic pain patients.

The ACC was initially thought to receive nociceptive information from the insular cortex (among other regions)^44,45^. However, neuronal tracing studies have shown that the claustrum rather than the insula provides a denser input to the ACC^46,47^. Consequently, the claustrum has only recently been investigated as a structure involved in pain processing^29,30^. Using both c-Fos expression and *in-vivo* calcium imaging, our results revealed a temporally dynamic response to noxious stimuli. We demonstrated that multiple modalities of nociception evoked a prolonged depression of activity that rebounded in the minutes after the stimulus. This rebound activity corresponded to a time frame when the animal is no longer displaying nocifensive behaviours and may relate to a period of resting state when nociceptive experiences are consolidated into longterm memory^24^. This premise is further supported by the observation that inhibiting claustrocingulate neurons impaired pain aversion in a conditioned pain aversion task (**Figure 2D**). Interestingly, while claustrocingulate lesion *impaired* nocifensive behaviors and pain aversion, it *enhanced* nociceptive sensitivity. This indicates that there is a disrupted relationship between reflexive behaviours and pain aversion, an observation that has been shown with other manipulations of ACC activity^18,48^. Overall, these results indicate that claustrocingulate neurons exhibit a multimodal response to nociceptive stimuli, and inhibition of this circuit can dissociate nociceptive and affective pain behaviours.

We also found that chronic inflammatory pain was associated with a loss of function in the claustrocingulate pathway. This observation is in accordance with the reduction in spontaneous claustrum activity and reduced synaptic communication at claustrocingulate synapses in chronic pain^29^. Loss of feedforward inhibition from the claustrum is a potential mechanism driving ACC hyperactivity associated with chronic pain. Accordingly, we found that chemogenetic activation of the claustrocingulate pathway reverses pain hypersensitivity, identifying a novel potential target for the management of chronic pain. Loss of claustrocortical activity may also contribute to cognitive impairment commonly observed in chronic pain^49^, although additional studies are needed to determine whether activation of this pathway can restore cognitive or affective pain behaviours.

The bimodal response of claustrocingulate neurons suggests this pathway subserves diverse and temporally-specific functions in processing nociceptive stimuli. Basal activity of claustrocingulate neurons may serve to filter nociceptive information in the ACC. Inhibition of this circuit, either through strong nociceptive stimuli, chronic pain, or chemogenetic manipulation, enhances nociceptive reflexive behaviours. This is in line with previous work that found reduced levels of claustrum activity are associated with increased impulsive motor action following a sensory cue^50,51^. The delayed claustrum activity that occurs well after nocifensive behaviours have resolved may serve a distinct function to coordinate a shift towards highly synchronous cortical states integral in quiet wakefulness and memory consolidation^24,52^. This bimodal conceptual framework could potentially resolve the seemingly conflicting theories on claustrum function ranging from sensory salience^50,53-56^ to resting state homeostasis ^24,52,57,58^.

This study identifies the claustrum as a central player in the pain neuromatrix. Claustrocingulate neurons display a multimodal response to nociceptive stimuli, and inhibition of this pathway dissociates the sensory and affective components of pain processing. Chronic pain results in a loss of function of this pathway and provides a novel mechanism to explain hypersensitivity and cognitive impairment associated with chronic pain. Future studies are needed to determine whether claustrocingulate neurons engage distinct neuronal ensembles to mediate sensory and affective pain behaviours. Validation of these results in other pain models is also required to ensure the consistency and broader applicability of our findings. Overall, this study reveals novel information for how the claustrum responds to and mediates pain and provides additional insight into the establishment and maintenance of chronic pain.

## 5. Acknowledgements

We would like to thank members of the Jackson lab and Taylor lab for comments on the manuscript, and the Faculty of Medicine & Dentistry Cell imaging Center including Xuejun Sun for help with microscopy. This work was supported by the Canadian Institutes of Health Research (Grant no. 426485 to JJ and 462798 to AMWT).

## 7. Figures

**Supplemental 1**. (A) Example images of PV (red) and c-Fos (green) co-labeling in saline and CFA treated mice following a noxious stimulus (scalebar = 100 µm). Across treatments we found no colocalization of PV and c-Fos. (B) Example images of SOM (red) and c-Fos (green) co-labeling in saline and CFA treated mice following a noxious stimulus (scalebar = 100 µm). Disparate colocalization was found but there was no difference between treatments (t(13) = 0.9748, p = 0.35). Data was analyzed by unpaired t-test. Data was collected from both male and female mice.

**Supplemental 2**. (A) CFA hind-paw injection significantly reduced paw withdrawal thresholds (t(13) = 3.897, p = 0.0018). The innocuous filament and noxious filament were chosen based on the withdrawal threshold of control mice. (B) Quantification of c-Fos+ cells not colocalized with CTB across treatments (H(4) = 36.10, p = < 0.0001). *p < 0.05, **p < 0.01, ***p < 0.001, ****p < 0.00001. Bars or scatterplots with error bars represent mean ± SEM. Data was analyzed by unpaired t-test or Kruskal-Wallis test with Sidak’s post hoc test. Data was collected in both male and female mice.

**Supplemental 3**. (A) Caspase lesion did not significantly reduce tail withdrawal latency (n = 11-15) (t(23) = 1.863, p = 0.0752). (B) CFA hindpaw injection reduced tail withdrawal latency in caspase mice compared to saline controls at 3 days post injection (n = 5-8) (effect of time: F(1.590, 30.22) = 2.961, p = 0.0775; effect of caspase: F(1, 19) = 10.11, p = 0.0049; effect of CFA: F(1, 19) = 7.990, p = 0.0108). (C) HM3Dq activation of claustrocingulate projections did not affect tail withdrawal latency, however there was a significant effect of CNO injection (n = 15-16) (effect of HM3Dq: F(1, 30) = 0.04187, p = 0.8392; effect of drug: F(1, 30) = 17.19, p = 0.0003). (D) CFA hindpaw injection did not reduce tail withdrawal latency up to 2 weeks post injection, HM3Dq activation also did not have an effect (n = 7-8)(effect of time: F(1.888, 52.85) = 0.6422, p = 0.5215; effect of caspase: F(1, 28) = 0.6173, p = 0.4386; effect of CFA: F(1, 28) = 0.5657, p = 0.4583). *p < 0.05. Data was analyzed by unpaired t test, or repeated measures ANOVA with Sidak’s post hoc test. Data was collected in male mice.

