## Supplemental Information for "Claustrum projections to the anterior cingulate modulate nociceptive and pain-associated behaviour"

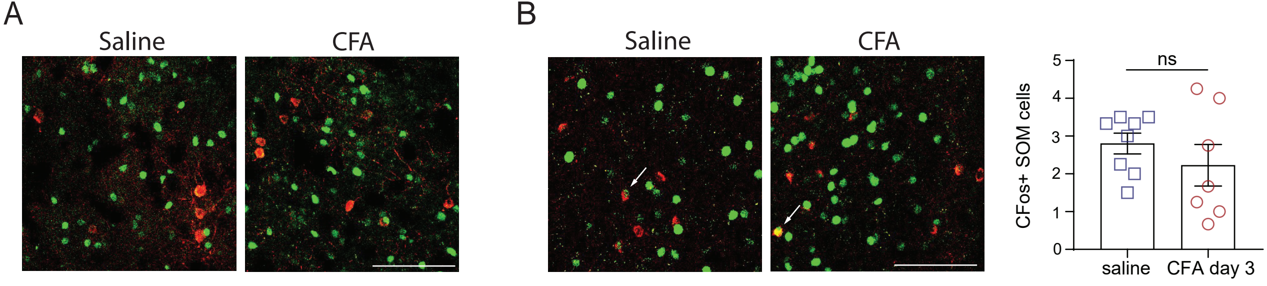
A
Saline
CFA
B
Saline
CFA
ns
CFos+ SOM cells
0
1
2
3
4
5
saline
CFA day 3

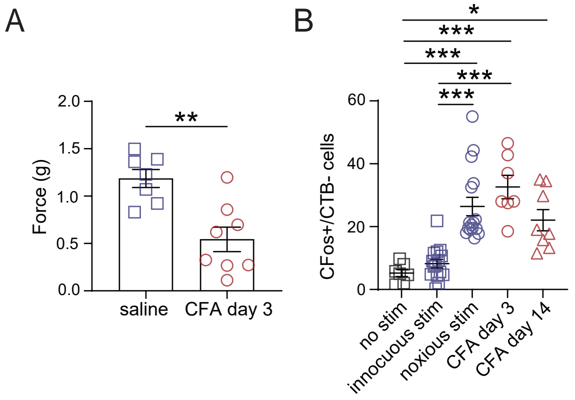
A
B
Force (g)
2.0
1.5
1.0
0.5
0.0
**
saline
CFA day 3
CFos+/CTB- cells
60
40
20
0
*
***
***
***
***
no stim
innocuous stim
noxious stim
CFA day 3
CFA day 14

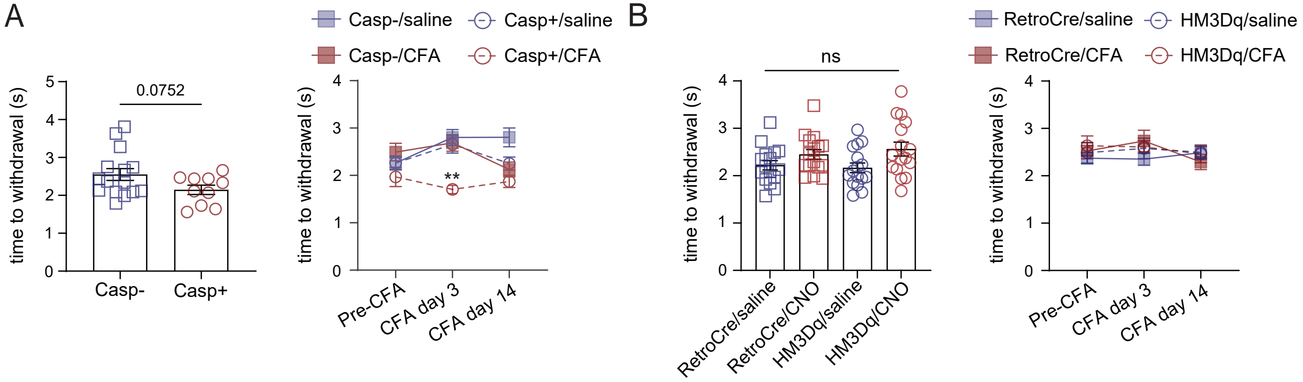
A
Casp-/saline
Casp+/saline
Casp-/CFA
Casp+/CFA
0.0752
time to withdrawal (s)
Casp-
Casp+
**
Pre-CFA
CFA day 3
CFA day 14
B
RetroCre/saline
HM3Dq/saline
RetroCre/CFA
HM3Dq/CFA
ns
RetroCre/saline
RetroCre/CNO
HM3Dq/saline
HM3Dq/CNO
